# Conservation and diversification of mating behavior patterns among three sibling species in the *Drosophila subobscura* species subgroup

**DOI:** 10.1101/2025.07.31.667830

**Authors:** Kenta Tomihara, Ryoya Tanaka, Francisco Rodríguez-Trelles, Daisuke Yamamoto

## Abstract

Behavioral traits are known to evolve rapidly, often even preceding morphological or physiological changes. However, the genomic and neural bases for such rapid behavioral changes remain to be clarified. *Drosophila subobscura* is a rare example of a species that performs nuptial gift giving, while the mating behaviors of the other two members of the *D. subobscura* species subgroup, *D. madeirensis* and *D. guanche*, remain largely unstudied. In the present study, we characterize and compare mating behaviors of three sibling species of the *D. subobscura* species subgroup, with the aim of providing a starting point for investigating the neural mechanisms underlying reproductive behavioral divergence in the D. *subobscura* subgroup. We find that *D. madeirensis* males exhibit a rich repertoire of courtship behaviors—very similar to that of *D. suboscura*—including tapping, midleg swinging, proboscis extension and nuptial gift giving. In contrast, *D. guanche* males perform only tapping and lack the other premating displays, yet they still copulate successfully. We postulate that female promiscuity has promoted the loss of multiple components of the male courtship repertoire in *D. guanche*. The relatively recent divergence among these species (∼1.72 Myr between *D. guanche* and other two species) suggests that only a few genomic and neural changes underpin the striking differences in mating behavior within the *subobscura* species subgroup. This system offers a promising platform for uncovering the mechanistic basis of rapid behavioral evolution.

## 1. Introduction

Animal behavior varies widely among species. Some closely related species are hardly distinguishable by morphologies, yet exhibit distinct behavioral repertoires. According to the “behavior evolves first” (BEF) hypothesis, behavioral changes precede and often influence the evolution of other traits, such as morphological and physiological traits (Rhodes & Kawecki, 2009). Although this idea has attracted much interest in evolutionary biology, supporting evidence remains scarce (Khan et al., 2024). In fact, it would be difficult to evaluate the validity of the BEF hypothesis without knowing what changes in genes and genomes underlie intraspecies behavioral differences and how such changes modify the neurons and neural circuits responsible for the behavior.

*Drosophila* provides an ideal platform for comparative studies of the genetic and neural bases of behavioral differences among species. Its suitability stems from the availability of genome sequence databases for many *Drosophila* species (Clark et al., 2007) and the complete brain connectome of the model species *D. melanogaster* (Dorkenwald et al., 2024; Schlegel et al., 2024), the latter of which can serve as a reference for identifying circuit-level differences underlying specific behaviors in other species.

We are interested in divergent mating behaviors, which are often shaped by species-specific sexual selection pressures (Debelle et al., 2014). Specifically, we focus on nuptial gift giving—a mating behavior in which males offer gifts to females to increase the likelihood of successful mating (Gwynne, 2008). Nuptial gift giving is widely found in the animal kingdom, from snails to humans (Lewis et al., 2011). In the genus *Drosophila*, nuptial gift giving has been reported in two species of the *obscura* group: *D. subobscura*, which belongs to the *D. subobscura* species subgroup (Immonen et al., 2009; Steele, 1986), and *D. persimillis*, a member of the *pseudoobscura* species subgroup (Hernández & Fabre, 2016). In the latter species, however, nuptial gift giving is not essential for successful mating. The neural circuit underlying nuptial gift giving in *D. subobscura* males includes neurons that express the male-specific transcription factor Fruitless M (FruM) (Tanaka et al., 2017). In both *D. melanogaster* and *D. subobscura*, FruM-expressing neurons form a courtship-related circuit known as the *fru* circuit. However, artificial activation of the *fru* circuit elicits different behavioral outcomes in the two species: nuptial gift giving is triggered in *D. subobscura* but not in *D. melanogaster*. This behavioral divergence appears to result from species-specific differences in circuit composition. In *D. subobscura*, insulin-like peptide producing cells (IPCs) express FruM and are integrated into the *fru* circuit, thereby playing a key role in orchestrating nuptial gift giving. In contrast, IPCs in *D. melanogaster* do not express FruM, are not part of the *fru* circuit, and do not contribute to courtship behavior (Tanaka et al., 2025). Thus, *D. subobscura* offers an unparalleled opportunity to explore the genomic and circuit-level mechanisms underlying the evolutionary emergence of novel, species-specific behaviors.

Whereas *D. subobscura* is a cosmopolitan species, two other members of the *subobscura* species subgroup are endemic species: *D. madeirensis*, a species endemic to the island of Madeira, and *D. guanche*, which is endemic to the Canary Islands (Figure 1). The three species are very closely related, and some combinations of crosses can even yield hybrids that are with or without fertility (Khadem & Krimbas, 1993; Krimbas & Loukas, 1984; Papaceit et al., 1991; Rego et al., 2006). In this study, we characterize courtship behaviors of the three species of the *D. subobscura* species subgroup with special reference to the presence or absence of nuptial gift giving, aiming to provide a starting point for future studies on the genomic and circuit bases that underlie the evolution of this distinct behavioral trait.

**Figure 1.**
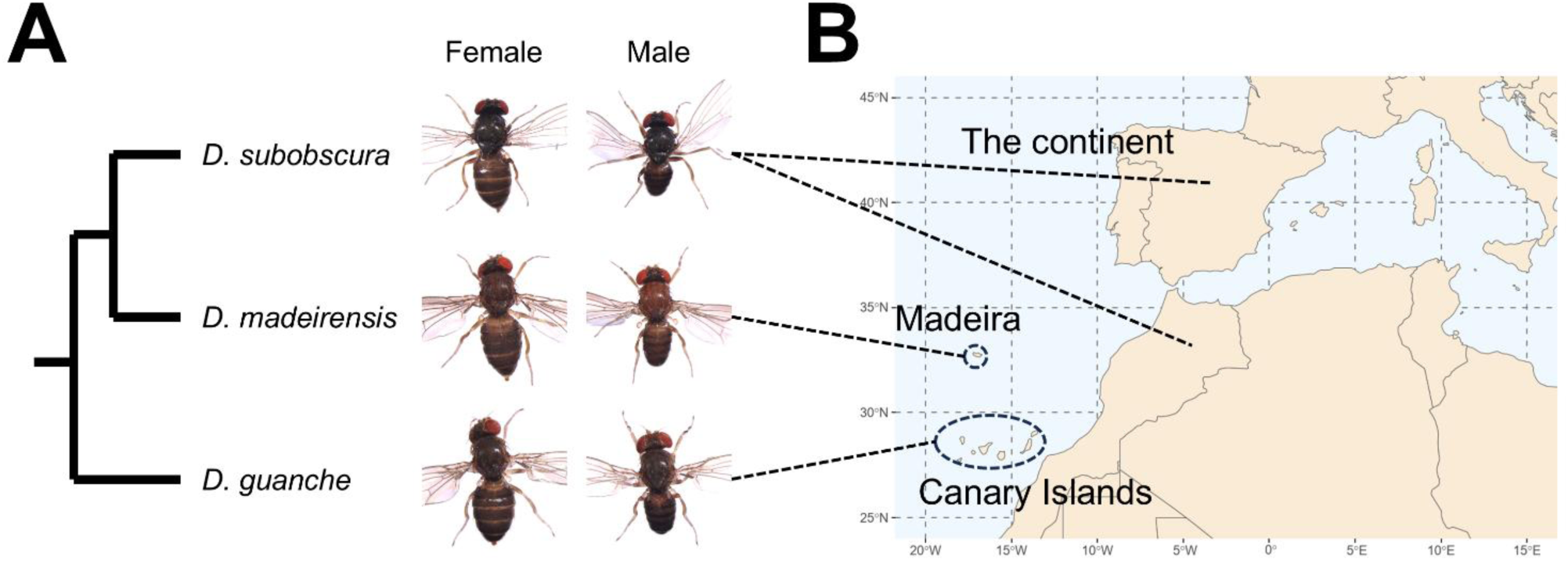
The three species comprising the *Drosophila subobscura* species subgroup. **(A)** Phylogeny of the *subobscura* species subgroup. (**B**) Geographical locations of the island of Madeira and the Canary Islands, which are inhabited by *D. madeirensis* and *D. guanche*, respectively.

## 2. Materials and Methods

### 2.1. Insects

*D. subobscura* strain 14011-0131.09 (collected in Combe Negre, France) was obtained from the Center for Insect Science, University of Arizona. *D. guanche* strain 14011-0095.01 (collected on the Canary Islands, Spain) was obtained from the San Diego Stock Center. *D. madeirensis* isofemale strain RF1 (a gift of Dr. Ana Llopart) was established from a single mated female collected in Ribeiro Frio, Madeira, Portugal (Herrig et al., 2014). Unless otherwise noted, insects were reared at 18°C on a cornmeal-yeast-agar medium and kept in an incubator under a 12/12 light-dark cycle.

### 2.2. Examination of the effect of light on mating

Adult flies in a vial were discarded and remaining pupae were kept under continuous light or dark conditions for 24 h. We then collected newly emerged flies from the vial, and 5–6 females were transferred to a single new vial with the same number of males, where they were allowed to mate freely under continuous light or dark conditions for 7 days. Each female was then transferred to a new vial, which was kept for approximately 2 weeks to check for the presence of offspring.

### 2.3. Examination of the remating rate

Newly emerged females were kept with and without males for 10–13 days to obtain virgin test females and mated test females, respectively. Males were single-housed for 10–13 days prior to the experiment. A pair of a single male and a mated or virgin female was placed in a chamber (φ40 mm × 6 mm) equipped with the video camera BFS-U3-32S4M-C (Teledyne FLIR, Wilsonville, OR, USA) or DFK 33UP1300 (The Imaging Source Asia, Taipei, Taiwan). Data acquired by video recording were subsequently analyzed to determine whether the female copulated with the male during the 1-h observation period. Throughout the experiment, the temperature was maintained at 21°C with an SCP-125 cooling plate (AS ONE, Osaka, Japan).

### 2.4. Quantification of elementary courtship actions

Virgin females were kept without males for 10–13 days and males were single-housed for 9–12 days prior to behavioral assays. To facilitate detection of the droplets of regurgitated crop contents on the labellum of courting males during video analysis, the males were fed red-colored sucrose solution (200 mM sucrose [Fujifilm Wako Pure Chemical, Osaka, Japan] and 2 μg/mL Food Red No. 106 [Tokyo Chemical Industry, Tokyo]) before behavioral assays. These males had been starved for 24 h beforehand to encourage them to drink the sucrose solution.

Single pairs of a male and virgin female were subjected to mating assays in a chamber (φ8 mm × 3 mm) for 15 min at 21°C under video recordings with the DFK 33UP1300 camera. The chamber used in this experiment was smaller than that used in the remating assay in order to increase the likelihood of an encounter. The courtship displays observed in a 5 min period starting 3 min after the introduction of flies into the chamber were manually annotated using ELAN software (Sloetjes & Wittenburg, 2008).

### 2.5. Examination of the structure of testes

Male flies of *D. subobsucra*, *D. madeirensis* and F_1_ hybrid were dissected at 5 days posteclosion, and testes were observed under an M205 FA microscope (Leica Microsystems, Wetzlar, Germany).

## 3. Results

### 3.1. Influence of light on the mating

Multiple studies have reported that *D. subobscura*, unlike most other *Drosophila* species, does not mate in the dark (Philip et al., 1944; Rendel, 1945; Ripfel & Becker, 1982; Wallace & Dobzhansky, 1946). The possible light dependence of mating in *D. subobscura*, *D. madeirensis* and *D. guanche* was examined by counting the females that yielded offspring when kept under dark conditions for 7 days with males (Figure 2(A)). We confirmed that no offspring were recovered in *D. subobscura* females when deprived of light. A comparable failure to produce offspring in darkness was seen in *D. madeirensis*, while *D. guanche* produced offspring under both light and dark conditions (Figure 2(B)).

**Figure 2.**
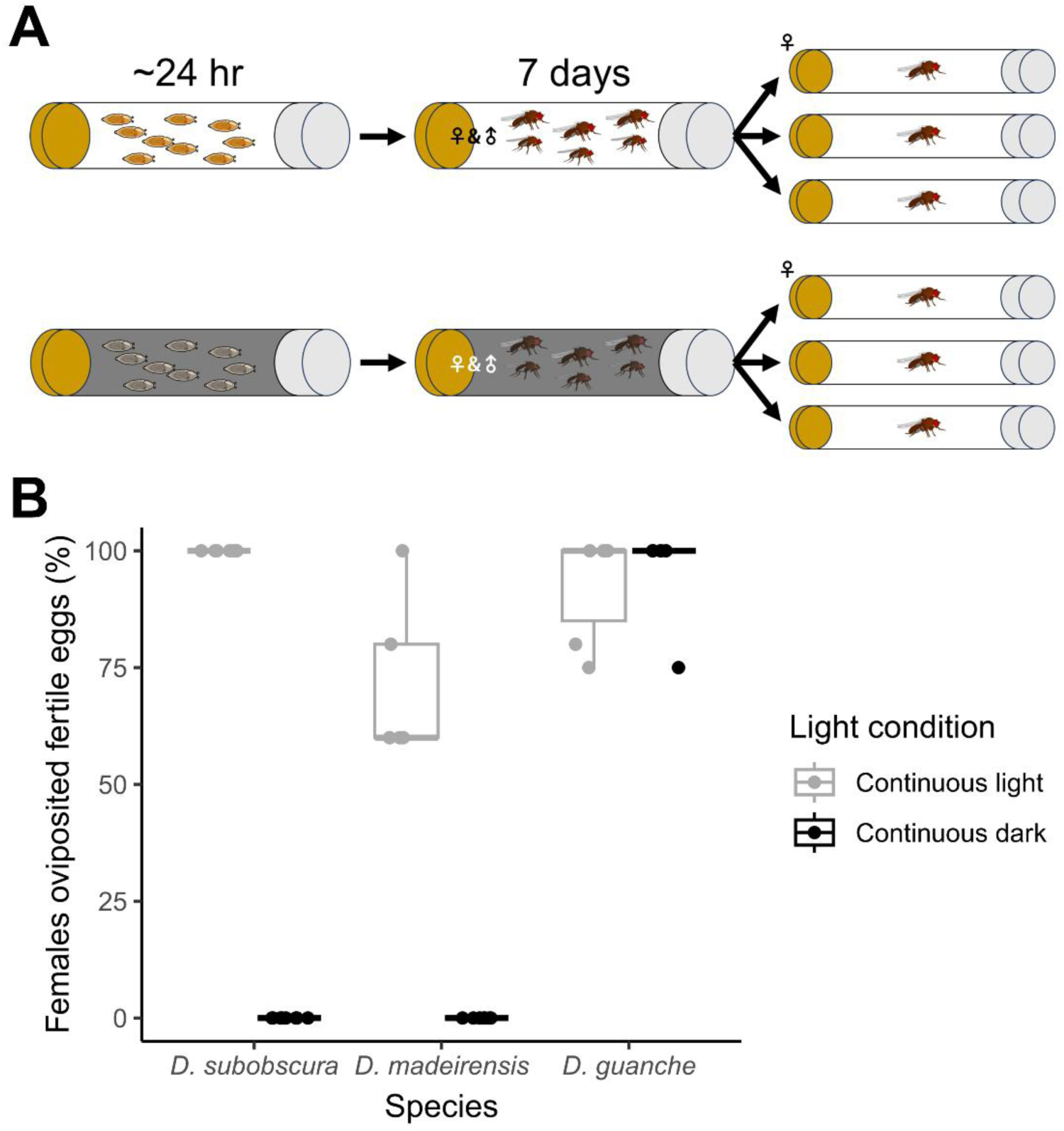
Comparisons of fertility among the three species of the *subobscura* species subgroup under light and dark conditions. (**A**) The experimental procedure. Approximately a day before eclosion, two test groups were established, one under a light and one under a dark condition. Upon eclosion, 5–6 virgin females and the same number of virgin males were transferred to a new vial for each test group and allowed to mate freely for 7 days under continuous light or dark conditions. Thereafter, single females were transferred to a new vial, and the presence or absence of offspring was scored. (**B**) The proportions of females that produced fertile eggs under the light (grey) and dark (black) conditions were scored for each of the three species. In the box plots, the upper quartile, median and lower quartile are indicated in addition to individual data points.

### 3.2 Remating

Monandry is a well-documented feature of *D. subobscura* females (Holman et al., 2008; Lizé et al., 2012; Maynard Smith, 1956), whereas *D. guanche* females have been reported to remate (Holman et al., 2008), suggesting a species-specific difference in female mating strategy. We confirmed these previous results by video-recording and subsequent analysis of mating events, in which a virgin or fertilized female was paired with a virgin male (Figure 3(A)). In *D. subobscura*, ∼50% of tested virgin females copulated, whereas none of the fertilized females did so (Figure 3(B)). In contrast, *D. guanche* pairs mated irrespective of whether the female was virgin or fertilized (Figure 3(B)). We further found that, in *D. madeirensis*, virgin, but not fertilized, females copulated under the current experimental conditions (Figure 3(B)). Our results were thus in line with the notion that *D. subobscura* females are monoandrous (Holman et al., 2008; Lizé et al., 2012; Maynard Smith, 1956). However, some previous studies have reported that a small proportion of *D. subobscura* females (∼8%) may remate, particularly if the first mating fails to produce offspring or if the ambient temperature exceeds ∼20°C (Fisher et al., 2013; Grandela et al., 2024; but see Loukas et al., 1981). We consider that monandrous mating in *D. subobscura* was preserved in the descendant species *D. madeirensis,* but was replaced by polyandrous mating in another descendant, *D. guanche*.

**Figure 3.**
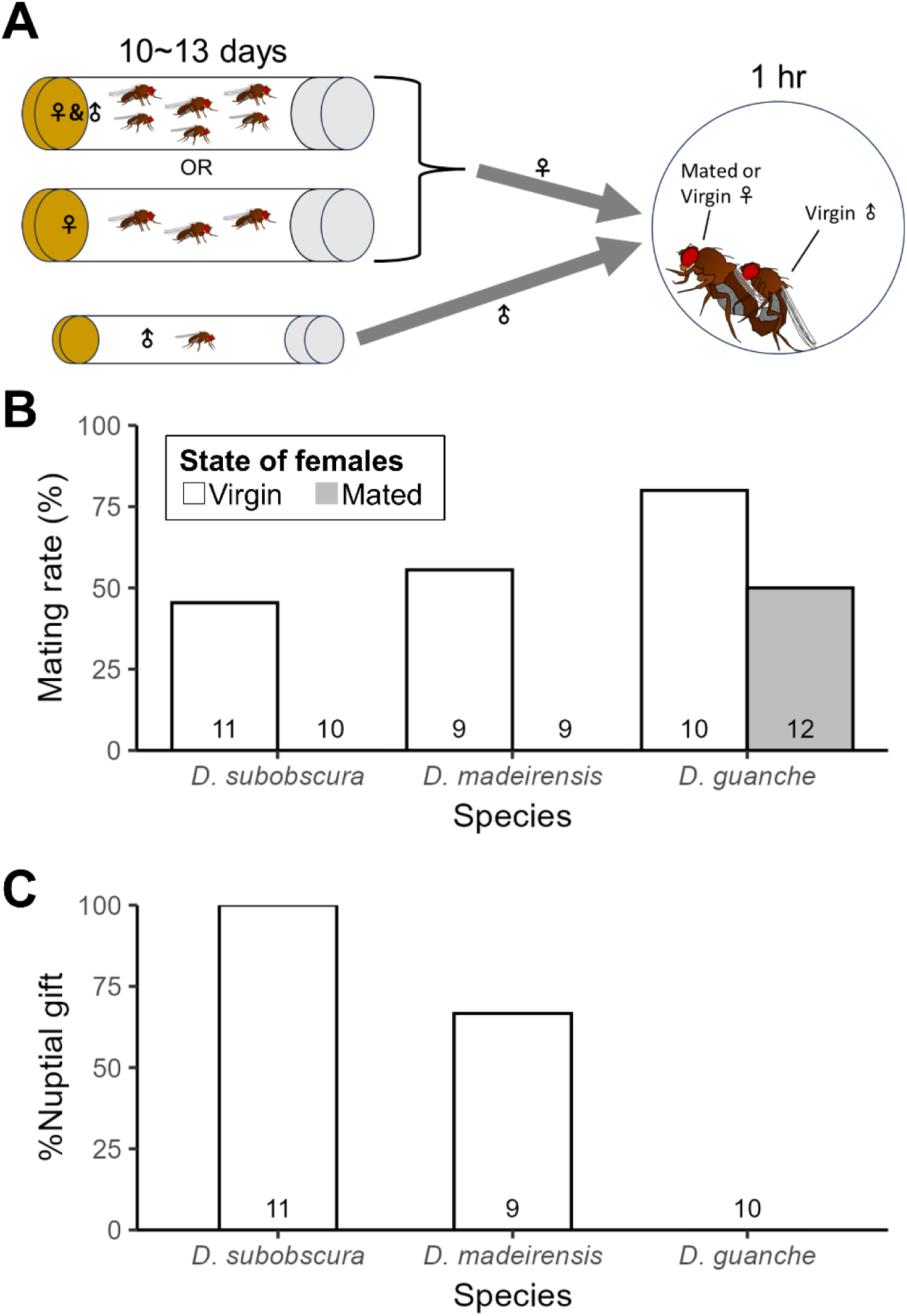
Comparison of remating rates and the prevalence of nuptial gift giving among the three species of the *subobscura* species subgroup. (**A**) The experimental procedure. Virgin and mated females were singly paired with a virgin male and their behavior was recorded with a video camera for 1 h to determine whether the females mated with the introduced male. (**B**) The proportion of virgin (open bars) and mated (hatched bars) females mated in a 1-h observation period. Note that only *D. guanche* remated under these conditions. (**C**) The proportion of males that showed nuptial gift giving within a 1-h observation period. All experiments were conducted in a large chamber of 40 mm in diameter and 6 mm in height.

### 3.3 Mating displays

We then studied how courtship proceeds by the analysis of mating behavior recorded in a video. To allow visual confirmation of gift-giving behavior, test males were fed a red-colored sucrose solution prior to mating assays. The color made regurgitated droplets on the labellum more easily detectable during video analysis (Figure 4(A)). In *D. subobscura*, six elementary steps that follow orientation have been described, i.e., tapping, midleg swing, proboscis extension with or without regurgitation, nuptial gift giving and attempted copulation, leading to copulation when successful (Higuchi et al., 2017; Spieth, 1952) (Figure 4(B), Video S1). Tapping refers to an action in which the male extends a foreleg and touches the body surface of a female. Midleg swing is characterized by bilateral posterior-to-anterior movements of a pair of the midlegs, typically repeated a few times in succession. Proboscis extension is performed when the male positions himself in front of a target female and may be repeated several times. Tapping, midleg swing and proboscis extension are commonly observed behavioral elements of a male when paired with a female. Only on limited occasions, the male may proceed to perform nuptial gift giving. In nuptial gift giving, a male fly presents regurgitated crop contents on the labellum protruded toward a female while extending two wings in a head-to-head position. Thereafter, the male rapidly turns around to get behind the female, then attempts to copulate, which is followed by copulation when successful. We found that *D. madeirensis* mating follows a behavioral sequence similar to that of *D. subobscura* (Figure 4(B-C), VideoS2). There was only one case of nuptial gift giving in *D. madeirensis* in the experiment shown in Figure 4(C), where a small chamber of 8 mm in diameter and 3 mm in height was used and the courtship displays observed in a 5 min period were scored. However, when a larger chamber of 40 mm in diameter and 6 mm in height was used and the observation time was extended to 1-h, more than half of tested *D. madeirensis* males engaged in nuptial gift giving (Figure 3(C), Video S2).

**Figure 4.**
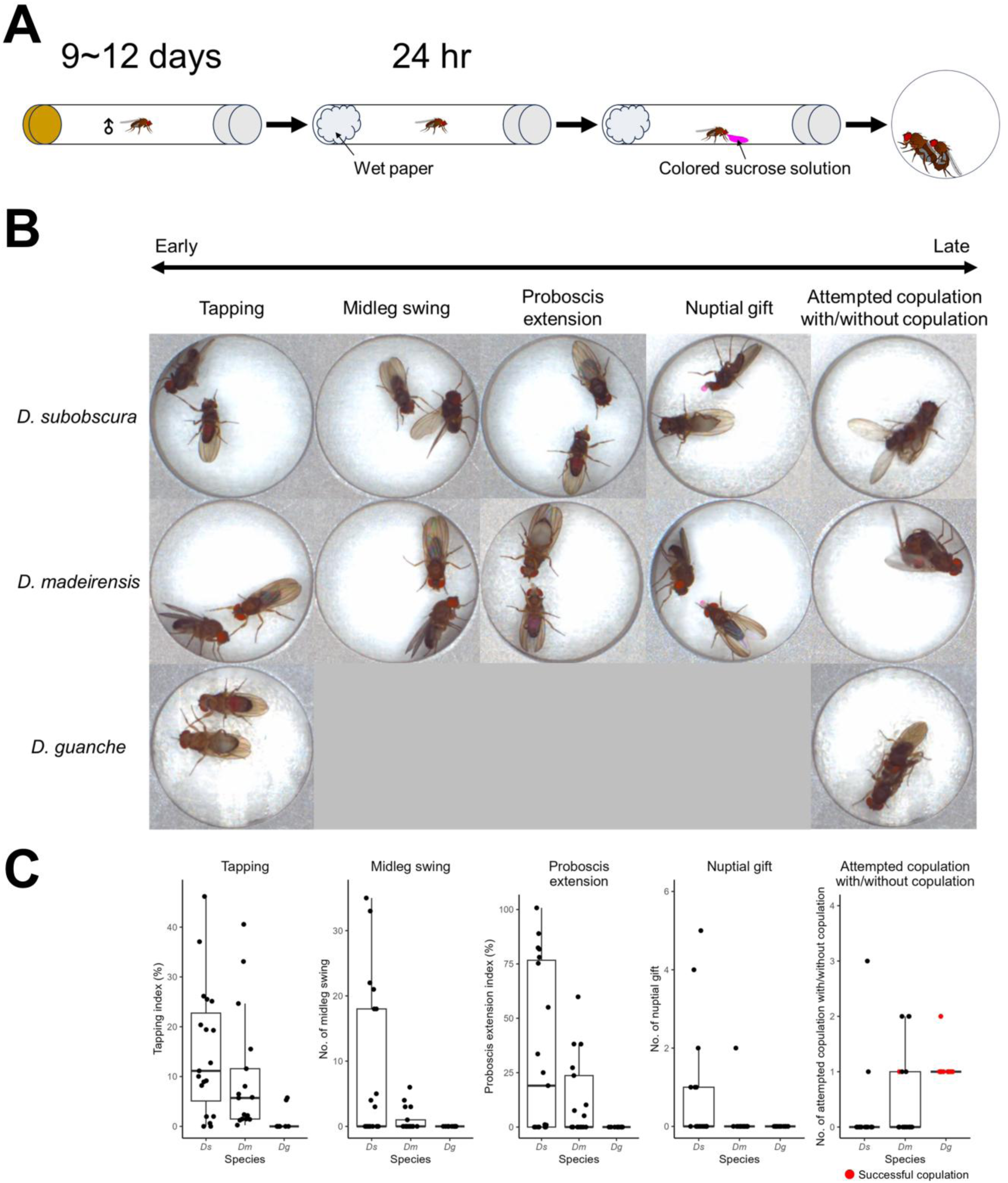
Comparison of mating behavior patterns among the three species of the *subobscura* species subgroup. (**A**) The experimental procedure. The test males kept singly after eclosion for 9–12 days in a vial containing food were transferred to a new vial that contained a water-immersed piece of paper but no food to starve test flies for 24 h. Thereafter, the males were allowed to take red-colored sucrose solution to facilitate the detection of regurgitated crop contents presented on the labellum during the video analysis. (**B**) Examples of major behavioral steps (tapping, midleg swing, proboscis extension, nuptial gift giving and attempted copulation with or without copulation, as indicated above each picture) were aligned according to the typical order of occurrence (in the order of early to late events, bottom). **(C)** Quantitative analysis of frequencies of five elementary actions. Mate pairs that successfully copulated within a timeframe are colored red. *Ds*: *D. subobscura*; *Dm*: *D. madeirensis*; *Dg*: *D. guanche*. A small chamber of 8 mm in diameter and 3 mm in height was used in the experiments shown in Figures 4, 5 and 6.

Mating behavior in *D. guanche* is strikingly different from that of *D. subobscura* and *D. madeirensis*. Once the *D. guanche* male found a female, he briefly tapped her body and immediately attempted to mount her without any discernible prior display: none of the actions of midleg swing, proboscis extension and nuptial gift giving were detected before the mounting action (Figure 4(B-C), Video S3). Thus, even though *D. guanche* is phylogenetically very close to *D. subobscura* and *D. madeirensis*, its mating behavior is drastically different from the latter two species.

### 3.4 Courtship in heterospecific pairs

When paired with a virgin female of *D. madeirensis* or *D. guanche*, *D. subobscura* males did not go beyond tapping in courting a female (Figure 5(A)). In the combination of a *D. subobscura* female and a *D. madeirensis* male, a few *D. madeirensis* males proceeded further, displaying midleg swing and proboscis extension (Figure 5(B)), although the frequency of these actions was lower in heterospecific courtship than in conspecific courtship (Figure 4(C)). In contrast, *D. madeirensis* males rarely went beyond tapping in courting a *D. guanche* female (Figure 5(B)). We conclude that males of both *D. subobscura* and *D. madeirensis* prefer conspecific over heterospecific females as courtship targets, indicative of premating isolation among the three species.

**Figure 5.**
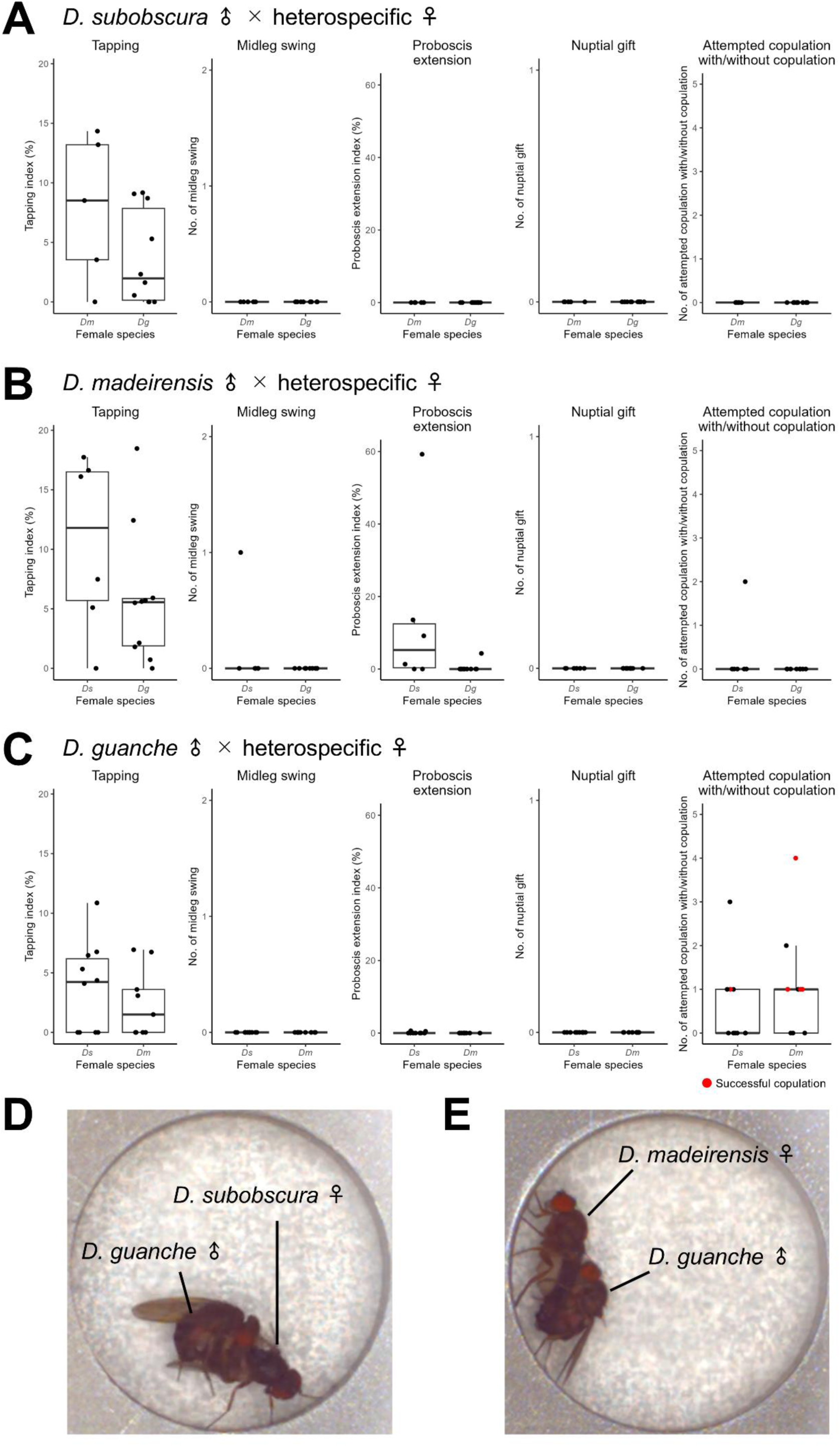
Courtship behaviors of *D. subobscura*, *D. madeirensis* and *D. guanche* males when paired with heterospecific females. (**A-C**) Quantitative analysis of five elementary actions in a male of *D. subobscura* (A), *D. madeirensis* (B), and *D. guanche* (C) paired with a heterospecific female. The elementary courtship actions analyzed include: tapping, midleg swing, proboscis extension, nuptial gift giving and attempted copulation with or without copulation. Pairs that successfully copulated within a timeframe are colored red. *Ds*: *D. subobscura*; *Dm*: *D. madeirensis*; *Dg*: *D. guanche*. (**D-E**) Examples of a *D. guanche* male courting a female of *D. subobscura* (D) and *D. madeirensis* (E).

When paired with virgin females of *D. subobscura* or *D. madeirensis*, the *D. guanche* males attempted to mount them and in some cases successfully copulated (Figure 5(C-E)), although the copulation rate in heterospecific courtship was lower than in conspecific courtship (Figure 4(C)). The heterospecific mating between *D. subobscura* or *D. madeirensis* females and *D. guanche* males did not yield an F_1_ hybrid in our experiment. Of note, (Krimbas & Loukas, 1984) succeeded in obtaining F_1_ hybrids between *D. madeirensis* females and *D. guanche* males, yet both sexes of F_1_ flies were sterile, demonstrating the postmating isolation between these species ((Krimbas & Loukas, 1984).

### 3.5 Courtship involving F_1_ hybrids between D. subobscura and D. madeirensis

Despite the high level of premating isolation, *D. subobscura* and *D. madeirensis* can produce hybrid offspring. We obtained F_1_ species hybrids from crosses between *D. madeirensis* females and *D. subobscura* males and examined the mating performance of F_1_ males when paired with a female of either parent species. Although F_1_ males were infertile due to the abnormal development of testes (Khadem & Krimbas, 1991) (Figure S1), they vigorously courted *D. subobscura* females (Figure 6, Video S4). The courtship behavior of F_1_ males followed an action sequence similar to that in *D. subobscura* and *D. madeirensis*: they displayed tapping, midleg swing, proboscis extension, and nuptial gift giving. In contrast, F_1_ males barely courted *D. madeirensis* females (Figure 6).

**Figure 6.**
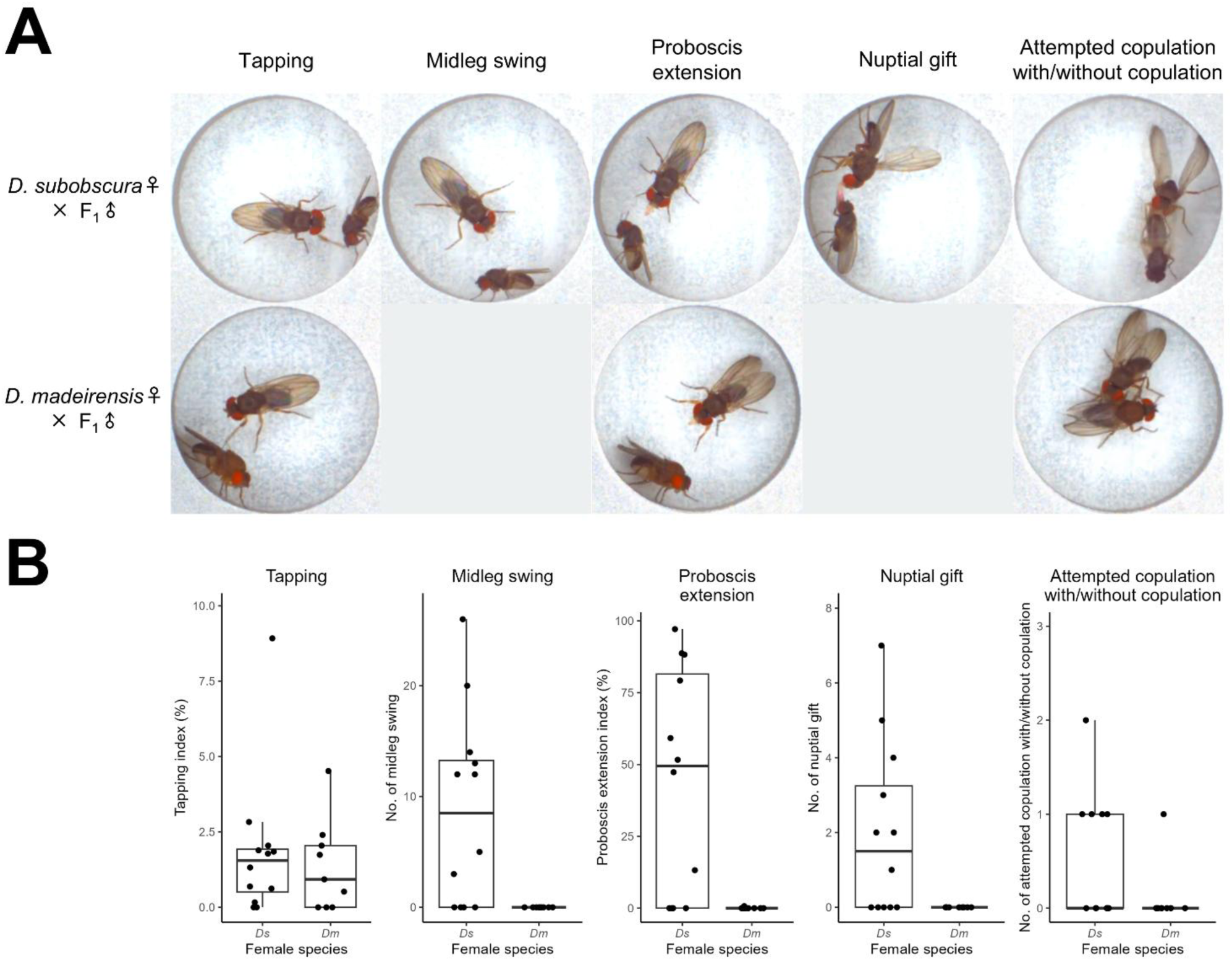
Courtship behavior patterns of hybrid males derived from a cross between *D. madeirensis* females and *D. subobscura* males. (**A**) Typical examples of courtship actions detected in a hybrid male courting a *D. subobscura* female (upper column) or a *D. madeirensis* female (lower column). (**B**) The ability of females to induce elementary courtship actions in hybrid males were compared between *D. subobscura* (*Ds*) and *D. madeirensis* (*Dm*). The elementary courtship actions analyzed include: tapping, midleg swing, proboscis extension, nuptial gift giving and attempted copulation with or without copulation. A single F_1_ male showed both proboscis extension and attempted copulation in courting a *D. madeirensis* female, while all other males exhibited no courtship action.

## 4. Discussion

This study revealed that, among the three sibling species composing the *subobscura* species subgroup, the mating behavior of *D. guanche* is markedly different from that of *D. subobscura* and *D. madeirensis*. Whereas males of *D. subobscura* and *D. madeirensis* display several discrete behavioral repertoires before taking an action of copulation, *D. guanche* males attempt to mount and copulate without any discernible prior displays except for brief tapping actions. In view of the fact that the deduced divergence time between *D. subobscura* and *D. madeirensis* is less than 1 million years (Myr) (Herrig et al., 2014), and that between *D. subobscura* and *D guanche* is 1.72 Myr (Karageorgiou et al., 2019), the new behavioral pattern must have arisen rapidly in *D. guanche* upon speciation. *D. guanche* is endemic to the Canary Islands, where this species likely gained the new behavior through the so-called “founder effect” (Carson, 1997; Mayr, 1942). It was predicted that when the population density is low on an island, less choosy females will have an advantage in obtaining a mating partner, because such females will require fewer courtship achievements by potential male partners before accepting them (Kaneshiro, 1976; but see Watanabe & Kawanishi, 1979). The relaxed requirement in courtship attempts will increase variabilities in behavioral displays by males, facilitating the development of new behavioral patterns. Interestingly, we found that *D. guanche* females repeat copulation. The observed high remating rate in fertilized females of *D. guanche* appears to represent attenuated sexual reluctance, which may have driven the rapid change in male mating behavior in this species.

Recently, we identified insulin-like peptide producing cells (IPCs) in the brain as a key element for executing nuptial gift giving in *D. subobscura* males (Tanaka et al., 2025). IPCs were shown to be postsynaptic to the *fru* circuit, which composes the core neural network for male courtship achievements (Tanaka et al., 2025). It is an open question whether the connection between IPCs and the *fru* circuit has been kept functional in *D. guanche*.

Immonene et al. (2009) hypothesized that *D. subobscura* males favor greater nutrition investment in nuptial gifts because, in this monandrous species, low levels of sperm competition ensure relatively secure paternity and reduce the risk of misdirected investment in unrelated offspring. Our finding that monandrous *D. subobscura* and *D. madeirensis* males provide nuptial gifts, while polyandrous *D. guanche* males do not, is consistent with the hypothesis proposed by Immonen et al. (2009). The males of *D. persimillis*, a polyandrous species belonging to the *pseudoobscura* species subgroup (Holman et al., 2008), also provide nuptial gifts, although this activity is not required for successful mating in this species (Hernández & Fabre, 2016). This might suggest that the regurgitation of crop contents during courtship was acquired in an ancestor common to the two related clades, i.e., the *subobscura* and *pseudoobscura* species subgroups, and subsequently established as nuptial gift giving in a monandrous ancestor of the two species, *D. subobscura* and *D. madeirensis*. Further studies of courtship behaviors of other members of the *obscura* species group—which includes both the *subobscura* and *pseudoobscura* subgroups—are needed to evaluate the hypothesis that nuptial gift giving by males and monoandry in females have coordinately evolved in these flies.

The present work showed that *D. subobscura* males more effectively discriminate conspecific from heterospecific females than do *D. madeirensis* (Figure 5(AB)). In the sibling species pair of *D. simulans* and *D. melanogaster*, premating isolation is maintained primarily by the differential preference for the sex pheromone 7,11-heptacosadiene (7,11-HD): 7,11-HD is a predominant female pheromone in *D. melanogaster* (Antony & Jallon, 1982) that attracts *D. melanogaster* males and repels *D. simulans* males (Billeter et al., 2009; Coleman et al., 2024; Sato & Yamamoto, 2020; Savarit et al., 1999). However, in the species of the *obscura* group examined by Khallaf et al. (2021), including *D. subobscura*, no female-specific pheromones were detected. Another sensory cue that often plays a key role in species recognition for mate choice in *Drosophila* is the production of species-specific courtship songs (Bennet-Clark & Ewing, 1969; Chen et al., 2019; Tootoonian et al., 2012). Although courtship songs in *Drosophila* are typically generated by wing vibration, courting males of *D. subobscura* do not vibrate their wings (Spieth, 1952). Absolute light dependence of successful mating may imply the involvement of vision, although possible visual cues for the species discrimination used by *D. subobscura* males escaped our detection. Thus, the sensory cues that allow *D. subobscura* males to distinguish conspecific from heterospecific females remain to be discovered.

Interestingly, we found that hybrid males obtained from the cross between *D. madeirensis* females and *D. subobscura* males vigorously court *D. subobscura* females but not *D. madeirensis* females (Figure 6), indicating that the allele inherited from *D. subobscura* acts dominantly over the one from *D. madeirensis* in the mating partner preference by the F_1_ hybrid males.

The whole genome sequence is available for all three members of the *subobscura* species subgroup (Karageorgiou et al., 2019; Puerma et al., 2018; Tomihara et al., 2024). What changes on the genome are responsible for the observed divergence in mating behavior among the three sibling species? This is another important question to be addressed in future studies. The three species yielded fertile hybrids when the cross was done in select directions (Khadem & Krimbas, 1993; Krimbas & Loukas, 1984; Papaceit et al., 1991; Rego et al., 2006). This opens the possibility of a forward genetic approach toward the identification of genes underlying behavioral diversification, in which introgression lines will be used to correlate unique behavioral traits to specific genetic rearrangements based on genomic sequence data.

## Acknowledgements

We thank Ana Llopart of the University of Iowa, for providing a *D. medeirensis* fly stock, and Manami Adachi for secretarial assistance. We also thank Toranosuke Nagamatsu for helpful discussions. The illustration of *Drosophila* pupa was acquired from TOGO TV (CC BY 4.0; https://togotv.dbcls.jp/).

## Conflict of Interest

The authors declare that they have no conflict of interest.

## Funder Information

This work was supported in part by a JSPS Grant-in-Aid (no. JP23KJ2210) to KT, a bioscience research grant of the Takeda Science Foundation to DY and a MEXT Grant-in-Aid (no. 25K21753) to DY.

## Author Contributions

KT, RT, and DY designed the study. KT conducted most of the experiments and analyses. FRT provided the insects used in this study. DY and KT drafted the manuscript. KT, RT, FRT, and DY reviewed and edited the manuscript. All authors approved the final version of the manuscript and agree to be accountable for all aspects of the work.

---

**Figure S1.**
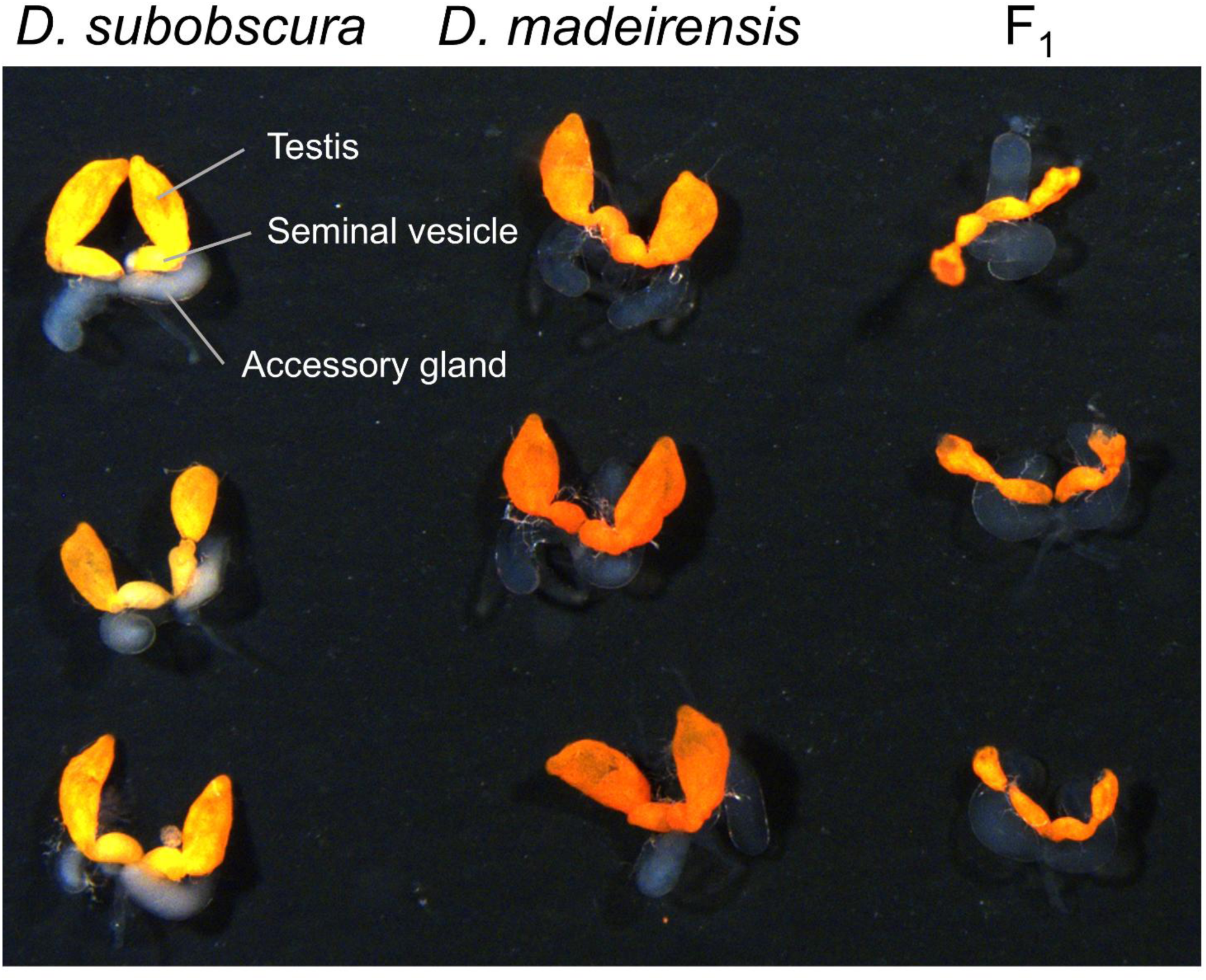
Testes from *D. subobscura*, *D. madeirensis*, and their F_1_ hybrid males. Three samples are shown for each species and hybrid. Hybrid males were obtained from a cross between *D. madeirensis* females and *D. subobscura* males.

**Supplemental video 1. Examples of courtship behaviors of a *D. subobscura* male courting a virgin *D. subobscura* female.**

**Supplemental video 2. Examples of courtship behaviors of a *D. madeirensis* male courting a virgin *D. madeirensis* female.**

**Supplemental video 3. Examples of mating behaviors of a *D. guanche* male courting a mated or virgin *D. guanche* female.**

**Supplemental video 4. Examples of courtship behaviors exhibited by a hybrid male derived from a cross between a *D. madeirensis* female and a *D. subobscura* male while courting a virgin *D. subobscura* female or a virgin *D. madeirensis* female.**

## Notes

### Competing Interest Statement

The authors have declared no competing interest.

